# Algorithms for Designing Therapeutic Oligonucleotides on Pangenome Graphs

**DOI:** 10.64898/2026.11.07.764582

**Authors:** Nathalie Gocht, Nicola Cotumaccio, Alexandru Tomescu, Alexander Schliep, Veli Mäkinen

## Abstract

Designing therapeutic oligonucleotides that act reliably across human genetic diversity requires more than alignment to a single reference genome. Binding affinity of an siRNA or ASO to a usually complementary site in an mRNA target molecule is one main determinant of drug efficacy. Thus, variation in sub-populations can lead to failures in drug development if this variation reduces binding affinity.

We provide a first methodological step towards principled, variant-aware design of therapeutic oligonucleotides. The genetic diversity in target genes, or specific regions thereof, is represented as a pangenome graph. We define individual or small sets of suitable therapeutic oligonucleotides by their ability to cover all paths in a directed acyclic graph within a bounded edit distance ≤ *k*. Edit distance is used as an accepted proxy to the commonly used binding affinity.

We formalize the resulting computational problem and provide complexity results: NP-hardness of identifying such minimal sets even for *k* = 0, and coNP-completeness^3^ of verifying the validity of a given set of oligonucleotides. Further, we propose algorithmic approaches operating on a product graph of an automaton (an Aho-Corasick DFA or an Ukkonen DFA) and the pangenome graph, as well as a dynamic programming-based lossless filter. We evaluate the approaches on gene-specific subgraphs of human pangenome graphs and demonstrate favorable performance compared to naïve brute-force computation.

Our results show that candidate target sites with a threshold of k ≤ 3 can be identified even for large, complex graphs where the brute-force computation did not complete within 24h. We observe that the dynamic programming-based lossless filter outperforms both product-graph approaches as well as bruteforce baselines while maintaining an average approximation ratio < 1.5 across more than a hundred evaluated subgraphs. We also present examples where a small set of oligonucleotides constitutes a feasible solution where no single oligonucleotide with a threshold of k ≤ 2 covers all paths.

## 1 Introduction

The algorithmic problems and solutions we present arise in the context of computational design of therapeutic oligonucleotides such as Anti-Sense Oligonucleotides (ASOs) [4] or small interfering RNA (siRNA) [8] when one wants to reflect a diverse genetic background represented as a pangenome graph in the design. State-of-the-art is evaluation of potential binding sites within a target gene sequence in a linear reference or patient genome. The therapeutic oligonucleotide candidates (our focus is on ASOs) are perfect or nearperfect Watson-Crick complements to the binding sites. Candidate ASOs are selected in multiple steps for desirable properties relating to drug efficacy and safety with algorithms, AI and experiments *in vitro* and *in vivo* and, finally, in clinical trials [4]. As the binding affinity between oligonucleotide and target mRNA is accepted as one of the main determinants of efficacy for ASOs [19, 32], variation in binding sites is clearly of concern. However, genetic variation in potential patients is typically only encountered late in the process, leading to a risk of expensive late-stage failure. As simplistic approaches, for example removing all regions with many Single Nucleotide Polymorphisms (SNPs) from consideration a priori, are likely to limit the choice of candidates too much, we investigate principled alternatives.

In particular, we focus on drug efficacy i.e. identifying, respectively confirming that, preferably one or a small number of therapeutic oligonucleotides will bind against a genetically diverse set of gene sequences represented as a pangenome graph. In order to focus on the elementary computational aspects, we abstract out many complex aspects related to oligonucleotide hybridization [2,4], and use sequence similarity, specifically edit distance, as a proxy. The algorithms we propose are a first important step towards integrating genetic variation early on in the *in silico* drug design process for therapeutic oligonucleotides. They help to prioritize high potential target sites in a systematic manner and potentially reduce the risk of late-stage failure.

Our approach targets a specific region of a pangenome graph rather than whole genes. Initially, such a region is partitioned into subgraphs (e.g. snarls [20]), that can be considered independently. We model the problem of finding a good set of oligonucleotide candidates for a given subgraph of a pangenome graph as follows. The set of oligonucleotides should be small, ideally of cardinality one, and each path (plausible haplotype) in our subgraph should be able to bind to one of the oligonucleotides. Figure 1 illustrates a case, where two strings are sufficient to cover all paths.

**Fig 1.**
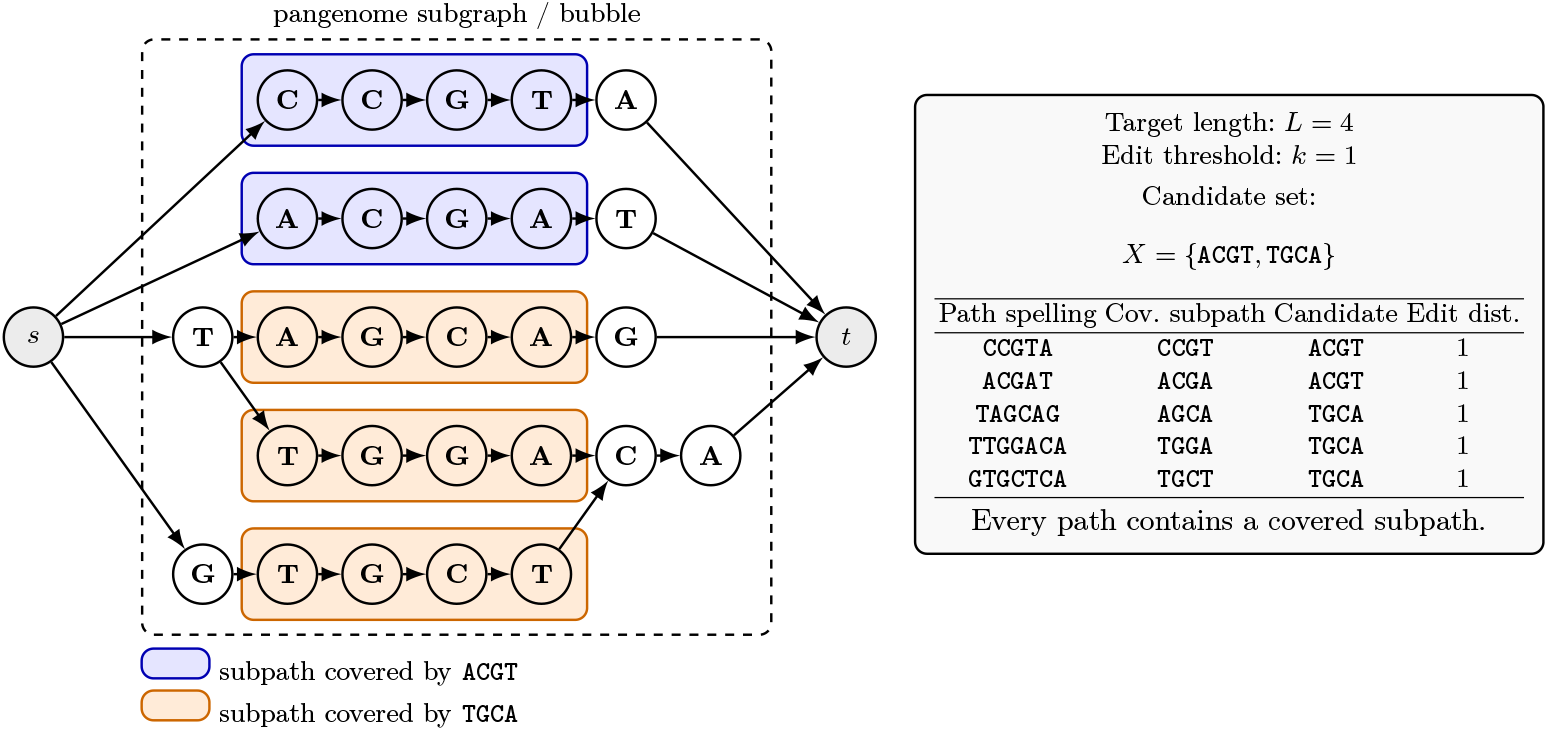
Illustration of the S<sc>tring</sc>-DAG-cover problem. Here, for clarity, source *s* and target *t* have no labels. The graph contains five source–sink paths spelling CCGTA, ACGAT, TAGCAG, TTGGACA, and `GTGCTCA`. For target length *L* = 4 and edit threshold *k* = 1, the candidate set *X* = {ACGT, TGCA} covers all paths. Importantly, the candidates need not match entire paths: each path only needs to contain some subpath within edit distance at most *k* from a candidate.

In the following, let *G* be a directed acyclic graph (DAG) with a unique source s and a unique sink *t* such that every node is labeled with a character from an alphabet *Σ*. We denote the set of states by *V*, the set of edges by *E* and the label of node *u* ∈ *V* by λ(*u*). The set of all finite strings on *Σ* is denoted by *Σ*\*. String *A* is a sequence of characters from *Σ* . Typically, we assume the characters in our alphabet are the nucleotides `A, C, G or T`. Edit distance *ed(A, B)* is the minimum number of single character substitutions, deletions, and insertions needed to convert string *A* into string *B* .

*Problem 1 (S<sc>tring</sc>-DAG-cover)*. Given a DAG *G*, a target length *L*, and an edit distance threshold *k*, find the minimum size of a set *X* ⊆ *Σ*^*^ of strings of length *L* such that each *s*-*t* path of *G* has a subpath spelling a string that has edit distance at most *k* from at least one string in *X* .

*Problem 2 (<sc>String</sc>-DAG-cover-verifying)*. Given a DAG *G*, a nonempty set *X* ⊆ *Σ*^*^ of strings, and an edit distance threshold *k*, decide whether each *s*-*t* path of *G* has a subpath spelling a string that has edit distance at most *k* from at least one string in *X* .

We show that Problem 1 is NP-hard with a reduction from set cover and that Problem 2 is coNP-complete for large *k* with a reduction from 3-SAT. We proceed to develop an approach based on the Aho–Corasick automaton and a graph product to solve Problem 2 when *k* = 0, that is, when no edits are allowed. Then we solve the general case by generating the edit neighborhood of the candidate strings, as well as the specific case of one string integrating Ukkonen automaton to the product graph approach. As these approaches are exponential in *k*, we also study a polynomial time filtering solution based on dynamic programming. We compare these approaches experimentally with the naive approach of extracting all paths and conducting approximate matching. We also enhance the framework with other aspects needed in oligonucleotide design, including targeting Problem 1 integrating an exhaustive enumeration of candidate strings. We additionally apply our approaches to the extension Problem 2 limiting the string to a fixed length *L*; since correctness for this setting has not yet been formally established, we report these results in the Appendix. Due to space constraints, also some proofs are deferred to the Appendix.

## 2 Method

### 2.1 Hardness results

Consider the *set cover problem* on a universe *U* = *{u*_*1*_,. .., *u*_*n*_}, where we are given *m* sets *S*_*j*_ ⊆ *U*, 1 ≤ *j* ≤ *m*, and we are asked to find the minimum collection of sets whose union is *U*. To cast this into Problem 1, it suffices to create a disjoint path from *s* to *t* for each element *u*_*i*_ ∈ *U* (in a relaxed setting where the DAG may have several sources and sinks; see the Appendix). For each *S*_*j*_ and for each element *u*_*i*_ ∈ *S*_*j*_ we add a node with label σ_*j*_ to the path corresponding to *u*_*i*_ in an arbitrary location. It then follows that a collection of sets whose union is *U* is in one-to-one correspondence with a set of strings of length *L* = 1 that cover all paths of the graph with edit distance *k* = 0.

This reduction can be further strengthened, so that the DAG labels come from an alphabet of size 3, instead of being *m* different labels (which also requires a different choice of *L*). Moreover, the DAG can be also structured so that all nodes lie on a single source-to-sink path. Additional edges from the end of an initial delimiter segment of this path, and into the start of a final delimiter segment, then model the source-to-sink paths from the initial reduction. The Appendix gives the complete proof.

#### Theorem3

*S<sc>tRing</sc>-DAG-COver. is NP-hard already for k* = 0, *even on DAGs whose nodes are labeled from an alphabet of size three, and whose nodes all lie on a single path (i*.*e. they consist of a single path, plus additional edges)*.

As we will soon see, the verification problem is easy for *k* = 0, but its complement problem turns out to be NP-complete for large *k*. This is shown by a reduction from 3-SAT. Recall that in 3-SAT we are to decide if a boolean formula consisting of clauses connected by and’s is satisfiable, where a clause consists of three variables or their negations (literals) connected by or’s. This reduction creates a DAG with *n* layers of two nodes each, one layer per variable, plus a source and a sink, where the source to sink paths correspond to all possible assignments of the *n* variables. Each clause maps to a string of length *n* in *X* containing the three falsifying literals and special symbols elsewhere. This string can be created so that a clause being false equals a path being covered with *k* = *n* − 3 edits, consisting of substituting the special symbols while matching the remaining 3 falsifying literals. Appendix gives the details of the reduction.

#### Theorem 4.

*S<sc>tring</sc>-DAG-cover-verifying is coNP-complete*.

#### Algorithm 1

The input consists of *G* and a DFA *A*= (*Q*, Σ, δ, *q*_*in*_,*F*) such thatℒ (*A*)= Σ^*\**^ ·*Y* . The output is “yes” iff every *s*-*t* path of *G* has a subpath spelling a string in *Y* .

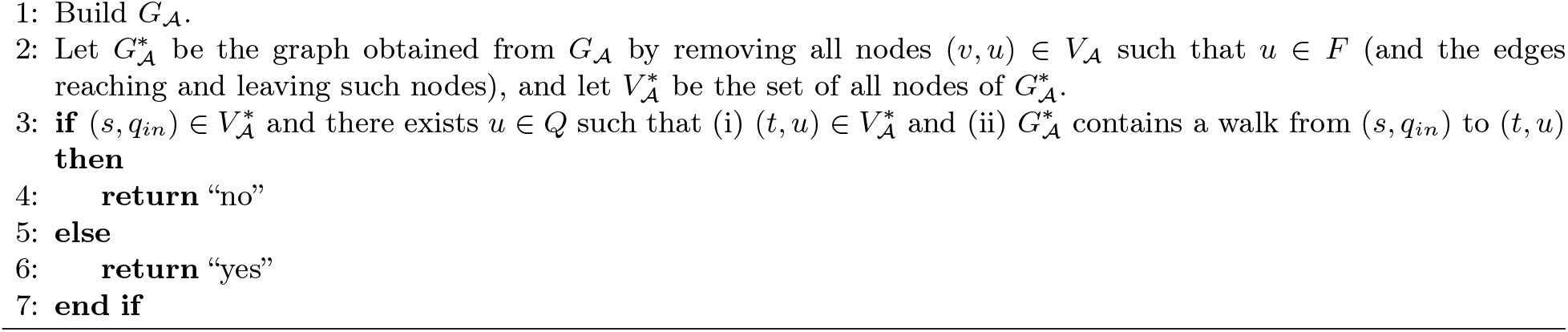

### 2.2 The product graph

To solve Problem 2 we can proceed as follows. Let *Y ⊆*Σ^*\**^ be a (possibly infinite) set, and consider the following problem: decide whether each *s*-*t* path of *G* has a subpath spelling a string in *Y* . Then, Problem 2 corresponds to the special case where *Y* is the set *N* ^*k*^(*X*) of all strings having edit distance at most *k* from at least one string in *X*. Consequently, we will first discuss how to solve the problem for an arbitrary *Y*, and then we will instantiate *Y* to *N* ^*k*^(*X*), distinguishing between the case *k* =0 and *k* arbitrary.

For technical reasons (required in the proof of Theorem 7 below), the sink *t* of *G* is not labeled with a character from Σ but with a new character $ *∉*Σ, i.e. λ(*t*) = $. Indeed, starting from *G*, we can create a new DAG *G*^*’*^ by adding a new sink *t*^*’*^ labeled with $ and then connecting the old sink *t* to the new sink *t*^*’*^. Then, each *s*-*t* path of *G* has a subpath spelling a string in *Y* iff each *s*-*t*^*’*^ path of *G*^*’*^ has a subpath spelling a string in *Y* (because no string in *Y* contains a character equal to $), so we can simply replace *G* with *G*^*’*^. Moreover, from now on we focus on the alphabet Σ = *{*A, C, G, T*}*.

We recall some basic notions from automata theory (see [7]). A (complete) deterministic finite automaton (DFA) is a 5-tuple *A* = (*Q*, Σ, δ, *q*_*in*_,*F*), where *Q* is a finite set of states, Σ is an alphabet, *δ* : *Q ×* Σ *→ Q* is the transition function, *q*_*in*_ ∈ *Q* is the initial state and *F ⊆ Q* is the set of final states. As usual in automata theory, *δ* is extended to the function from *Q ×* Σ^*\**^ to *Q* such that, for every *q ∈ Q* and for every *x ∈* Σ^*\**^, *δ*(*q, z*) is the state that we reach after reading *z* on *A* starting from *q*. The language *ℒ*(*A*) recognized by *A* is the set of all strings *z ∈* Σ^*\**^ such that *δ*(*q*_*in*_, *z*) *∈ F*.

In general, given a DFA *A*, we can define a *product graph* between *G* and *A*. The idea is that we can use a product graph to synchronize the paths in *G* and the runs of *A* on different input strings.

#### Definition 5.

*Let A*= (*Q*, Σ, δ, *q*_*in*_,*F*) *be a DFA. The product graph G*_*A*_ *is the (directed, unlabeled) graph defined as follows:*

- *The set V*_*A*_ *of nodes is V × Q*.
*The set E*_*A*_ *of edges in the given by all pairs* ((*v*_1_, *q*_1_), (*v*_2_, *q*_2_)) *∈V*_*A*_*× V*_*A*_ *such that* (*v*_1_, *v*_2_) *∈ E and q*_2_ = *δ*(*q*_1_, λ(*v*_1_)).

#### Remark 6.

Notice that |*E*_*A*_| = |*E*| *·*|*Q*| . Indeed, the function *ϕ* : *E ×Q →E*_*A*_ such that ((*v*_1_, *v*_2_), *q*_1_) ↦ ((*v*_1_, *q*_1_), (*v*_2_, δ(*q*_1_, λ(*v*_1_)))) is a bijection.

Let us show that, to decide whether each *s*-*t* path of *G* has a subpath spelling a string in *Y*, we only need to consider a DFA that recognizes all strings suffixed by some string in *U*. More precisely, we need a DFA *A* such that *ℒ* (*A*)= Σ^*\**^ *· Y* (where *·* is the standard concatenation of languages).

#### Theorem 7.

*Let Y ⊆* Σ^*\**^ *be a set of strings, and let A* = (*Q*, Σ, δ, *q*_*in*_,*F*) *be a DFA such that ℒ* (*A*)= Σ^*\**^ *·Y* .

*Then, Algorithm1 returns “yes” iff every s-t path of G has a subpath spelling a string in Y* .

*Proof*. See Appendix B.1.

The running time of Algorithm 1 is linear in the size of the product graph, as shown next.

#### Lemma 8.

*Given G and the DFA A, Algorithm 1 can be implemented in O*(|*E*|*·* |*Q*|) *time. Proof*. See Appendix B.2.

We now focus on Problem 2. To apply Theorem 7, we need to build a DFA *A* such that *ℒ* (*A*)= Σ^*\**^*·N* ^*k*^(*X*).

### 2.3 The case *k* = 0

Let us consider the case *k* =0 in Problem 2. Then, we only allow exact matches, and *N* ^*k*^(*X*)= *X*. We need a DFA *A* such that *L*(*A*)= *Σ*^*\**^ · *X* and, to this end, we can resort to the *Aho–Corasick* automaton for *X* [1], thus following an approach similar to the one proposed in [28] (where an Aho–Corasick automaton is used to solve the *shortest path problem with forbidden paths*).

#### Definition 9.

*( [1]) Let X ⊆ Σ*^*\**^ *be a nonempty set of strings. The Aho–Corasick automaton A*_*X*_ *of X is the DFA A*_*X*_ = (*Q, Σ, δ, q*_*in*_,*F* ) *defined as follows:*

- *Q is the set* Pref(*X*) *of all strings that are a prefix of at least one string in X. Note that the empty string belongs to Q*.
- *For every q ∈ Q and for every a ∈ Σ, δ* (*q, a*) *is the longest suffix of qa that belongs to* Pref(*X*).
- *q*_*in*_ *is the empty string*.
- *F is the set of all q ∈ Q such that at least one string in X is a suffix of q*.

Aho–Corasick automata satisfy the following crucial property, which intuitively follows from the definition of the transition function.

#### Lemma 10

**( [1])**. *Let ⊆ Σ*^*\**^ *be a nonempty set of strings, and let A*_*X*_ = (*Q, Σ, δ, q*_*in*_,*F*) *be the Aho– Corasick automaton of X. Fix z ∈ Σ* ^*\**^. *Then, δ* (*q*_*in*_, *z*) *is the longest suffix of z that belongs to* Pref(*X*).

Lemma 10 implies that *A*_*X*_ has the property required to apply Theorem 7.

#### Corollary 11

**( [1])**. *Let X ⊆ Σ*^*\**^ *be a nonempty set of strings. Then, L*(*A*_*X*_ )= *Σ*^*\**^ *· X*.

Moreover, *A*_*X*_ can be built in linear time.

#### Lemma 12

**( [1])**. *Let ⊆ Σ*^*\**^ *be a nonempty set of strings. Then, A*_*X*_ *can be built in O*(*Σ*_*x∈X*_ |*x*|) *time*.

We can solve Problem 2 for *k* = 0.

#### Corollary 13.

*Let X ⊆ Σ*^*\**^ *be a nonempty set of strings. In O*( |*E*|*Σ*_*x∈X*_ |*x*|) *time, we can decide whether each s-t path of G has a subpath spelling a string in X*.

*Proof*. See Appendix B.3.

### 2.4 Arbitrary

We now discuss Problem 2 when *k* is arbitrary. We focus on the case where *X* contains a single string *x* ∈ *Σ*^*^, so we write *N*^*k*^*(x)* instead of *N*^*k*^*(X*) (for the case |X| *>* 1, see Section 4). To apply Theorem 7, we need to build a DFA *A* such that ℒ (*A*) = *Σ*^*^ • *N*^*k*^ (*x*). While it is possible to obtain such a DFA by generating *N*^*k*^ *(x)* and then building the Aho-Corasick automaton of *N*^*k*^ *(x)*, Ukkonen showed how to build a DFA tailored for our problem [27].

By definition, we have *N*^*k*^*(x)* = *{z* ∈ Σ^*\**^ | *ed(x, z)* ≤*k*}. The set Σ^*^ • *N*^*k*^*(x)* can be characterized as follows. If *z* = *b*_*1*_*b*_*2*_ .. .*b*_*n*_, define *ed*^*\**^*(x,z)* = min_1≤*ℓ* ≤*n*+1_ *ed(x,b*_*ℓ*_*b*_*ℓ*_*+1* .. .*b*_*n*_). Then, *Σ*^*\**^ • *N*^*k*^*(x)* = *{z* ∈ *Σ*^*\**^ | *ed*^*\**^(*x, z*) ≤ *k}*.

The expression *ed*^*\**^ (*x, z*) is a well-known variant of the edit distance, classically used to solve the approximate pattern matching problem [24]. Algorithmically, the quantities *ed(x, z)* and *ed*^*\**^*(x, z)* can be computed using the same dynamic programming approach. More precisely, let *x* = *a*_*1*_*a*_*2*_ . .. *a*_*m*_ and *z* = *b*_*1*_*b*_*2*_ .. . *b*_*n*_, and consider a matrix *D* = *(d*_*ij*_)_*o≤i≤m,o≤j≤n*_. For 1 ≤ *i* ≤ *m* and 1 ≤ *j* ≤ n, define recursively:

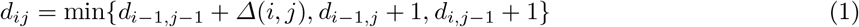

where Δ *(i,j)* = 0 if *ai* = *bj* and Δ *(i,j)* = 1 if *ai* ≠*bj*. Then, *D* is uniquely determined once we define *dij* for *(i* = 0) V *(j* = 0). Let *d*_*i0*_ = *i* for 0 ≤*i* ≤ m. Then:

**—** If we define *d*_*0j*_ = *j* for 0 ≤ *j* ≤*n*, we obtain *d*_*mn*_ = *ed(x, z)* [18, 29].

If we define *d*_0*j*_ = 0 for 0 ≤ *j* ≤ n, we obtain *d*_*mn*_ = *ed*(x, z)* [24,26,27].

To solve Problem 2, we will use the *Ukkonen DFA* for *x* [27]. Ukkonen defines a DFA using an algorithmic procedure. We now introduce the same DFA via a self-contained definition.

#### Definition 14

**( [27])**. *Let x* = *a*_*i*_ *a*_2_ …*a*_*m*_ ∈ *∑* and let k* ≥ 0. *For every z = b*_*1*_*b*_*2*_ …*b*_*n*_ ∈ *∑*, let* 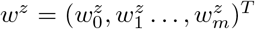 *the vector defined by* 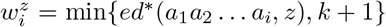 *for every* 0 ≤ *i* ≤ *m. The Ukkonen DFA* 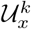 *of x is the DFA* 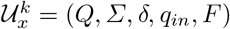 *defined as follows:*

- *Q = {w*^*z*^ | *z* ∈ *∑*}*.
- *For every z* ∈ *∑* and for every b* ∈ *∑, δ (w*^*z*^, *a)* = *w*^*za*^.
- *q*_*in*_ = *w*^*∈*^, *where ∈ is the empty string*.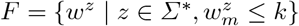
- *F is the set of all q ∈ Q such that at least one string in X is a suffix of q*

Every state of 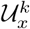is a vector of length m. We may also define 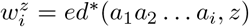, but Ukkonen shows that we can reduce the number of states by defining 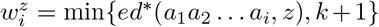[27]. The intuition is that 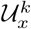only need to store the values of *ed\** up to *k* and remember which values are larger than *k, so we* use *k* +1 to refer to any value larger than *k*. Definition 14 is strictly related to the matrix *D* = *(d*_*ij*_)_*1*_*≤i≤*_*m*_,_*1*_*≤j≤*_*n*_ define before. More precisely, if we fix *z* = *b*_*1*_, *b*_*2*_, .. .,*b*_*n*_ and we define 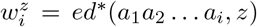, then 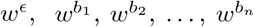 are the columns of *D* listed from left to right (under the initial condition *d*_*0j*_ = 0 for 0 ≤ *j* ≤ n). In particular, Equation 1 ensures that in Definition 14 the definition of *S* does not depend on the choice of z, that is, whenever 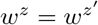, we have 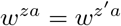. Lastly, the definition of *F* implies that, for every *z* = *b*_1_, *b*_2_,…, *b*_n_, we have 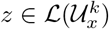 iff *ed*(x, z)* < t. Since *Σ\** . *N*^*k*^(x) = *{z* ∈ *∑*\* | *ed\** (x, z) ≤ k}, we intuitively obtain the following corollary (and so we can apply Theorem 7).

**Corollary 15 (** |**27**|). *Let x* ∈ *∑* and let k* ≥ 0. *Then*, 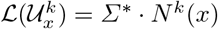.

For fixed *k* and constant alphabet, Ukkonen’s paper yields the following bounds.

#### Lemma 16

**( [27|)**. *Let x* ∈ *∑* and let k* ≥ 0. *Then*, 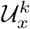*has O*(|*x*|^k+1^) *states and can be built in O*(|*x*|^k+2^) *time*.

Notice that, for *k* = 0, we have a linear number of states, analogously to the Aho-Corasick automaton of x. However, for *k* = 0, the construction time is quadratic. Intuitively, for every *k* > 0 the construction time is *O*(|*x*|^k+2^) and not *O*(|*x*|^k+1^) because every state is a length of vector |*x*| and so we need *O*(|*x*|) time per state. However, when we store 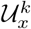, we do not represent every state as a vector, but we map distinct vectors to distinct integers (Ukkonen follows the same approach [27]). By doing so, after construction, each state and each transition can be processed in constant time. In particular, the time bound in Lemma 8 holds when the input DFA is 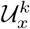.

We can finally show how to solve Problem 2.

**Corollary 17**. *Let x* ∈ *∑* be a string, and let k* > *0. In O(*l*E*|-|*x*|^k+1^ + |*x*|^k+2^) *time, we can decide whether each s-t path of G has a subpath spel ling a string that has edit distance at most k from x*.

*Proof*. See Appendix B.4.

### 2.5 Lossless filter for String-DAG-Cover-Verifying through dynamic programming

While we know that Problem 2 is coNP-complete, there is hope that it can be solved in some family of graphs. Towards such characterization, we develop a dynamic programming approach that works as a *lossless filter (LLF)* : it solves the problem correctly in many cases, but can fail to report correct solutions in some cases. We explain the approach with one candidate string *x* ∈ *X* and then extend the approach to the general case |XI *>* 1.

For a graph *G* as defined in Section 1 with a single source node, *s*, and a single sink node, *t*, we aim to compute the values *D*_*x*_*(v, i)* defined as the minimum edit distance between prefix x[1..i] and suffix of a worst path ending at node *v*:

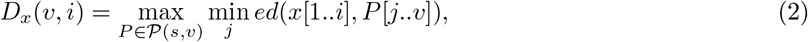

where *P*(*s,v*) denotes the set of strings spelled by paths from *s* to *v*.

We *estimate* these values using the following recurrence:

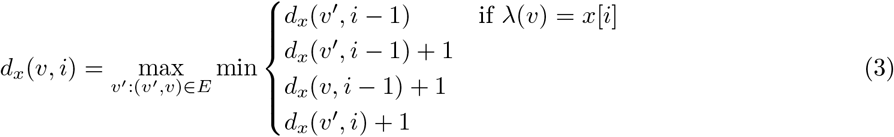

After linear-time topological sort of *G* and after initializing *d(v*, 0) = 0 for all *v* to allow alignments to start anywhere, we can compute all values *d*_x_(*v,i*) in time O(|E| · L).

Now let *c*_*x*_(*v*) = *D*_*x*_(*v, L*), that is, the number of edits needed to align *x*[1..L] to the worst possible path ending at *v*. We aim to compute the bottleneck cost *B*_*x*_ (*t*), defined as the cost of aligning *x* on the worst possible path from *s* to *t*. We estimate each c_*x*_(*v*) = *d*_*x*_*(v, L)* and use the following recurrence to estimate values *B*_*x*_(*v*):

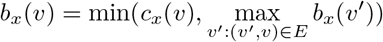

After linear-time topological sorting of *G*, we can compute *b*_*x*_(*t*) in *O*(|E|) time. Now, if *b*_*x*_(*t*) < *k* then *X* = {*x*} is a feasible solution to our problem.

The filtering approach can be extended to the general case |*X* | > 1 by conducting the first pass of dynamic programming independently for each string and taking the minimum of these values during the second pass of dynamic programming. Figure 2 illustrates this computation on an input, where the approach gives the correct solution.

**Fig 2.**
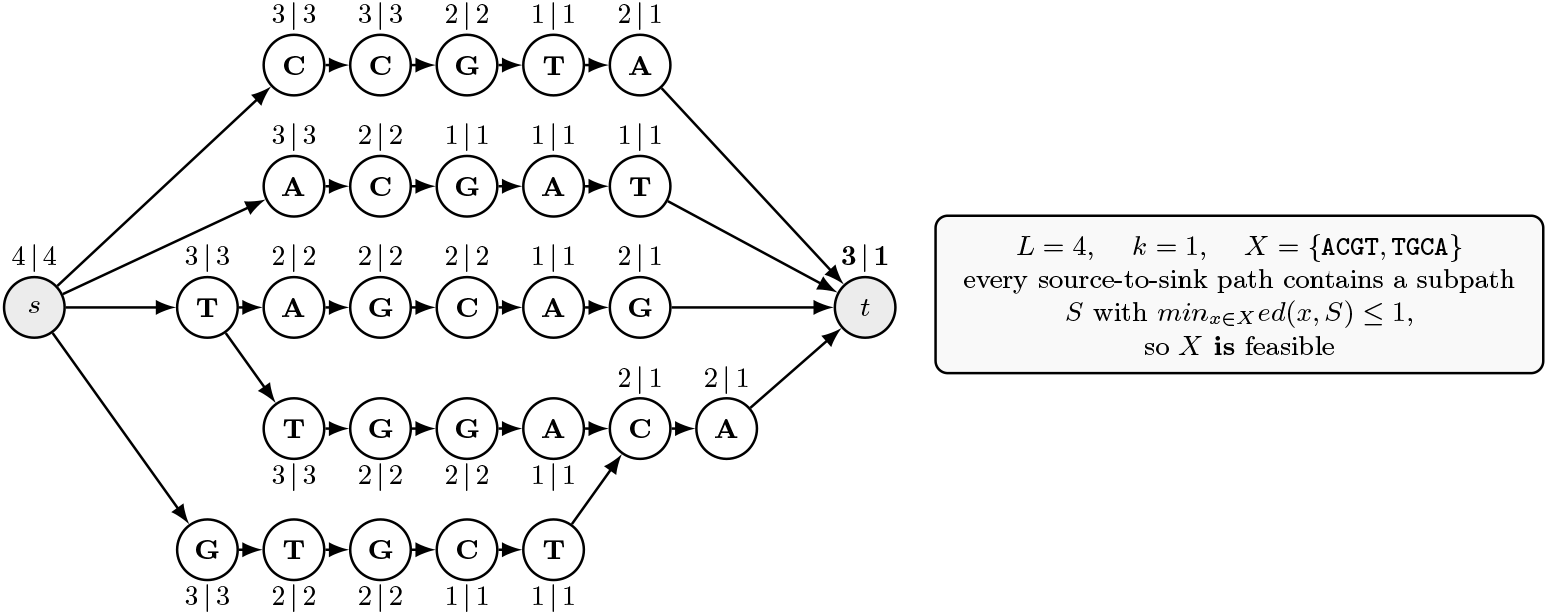
The two-pass dynamic programming scheme on the instance of Figure 1. Each node *v* is annotated *c*_*X*_ (v) | *b*_*X*_ (v). *First pass: c*_*X*_ (v) is the dynamic programming estimate on the cost min*x*_∈ *X*_ *D*_*x*_ (v, L) of aligning the best candidate in *X* to the *worst* source-to-*v* path, minimised over the subpaths S ending at *v* — that is, the bottleneck cost accumulated at *v*. In this example, the dynamic programming estimation gives the correct result for each node *v. Second pass: b*_*x*_(v) = min*(c*_*x*_(v), max_*u*:_(_*u*_,_*v*_)_∈_*E b*_*x*_*(u)*, which propagates the best coverage already achieved upstream, taking the maximum over predecessors so that *every* incoming path must have been covered. Since *b*_*X*_ *(t)* = 1≤k, the scheme certifies that every source-to-sink path is covered.

It is easy to find counterexamples for both passes of this dynamic programming approach. For example, already the first pass fails on the reduction graph of Theorem 4. Figure 3 illustrates a real pangenome subgraph found in our experiments, where the first pass is correct, but the second pass fails.

**Fig 3.**
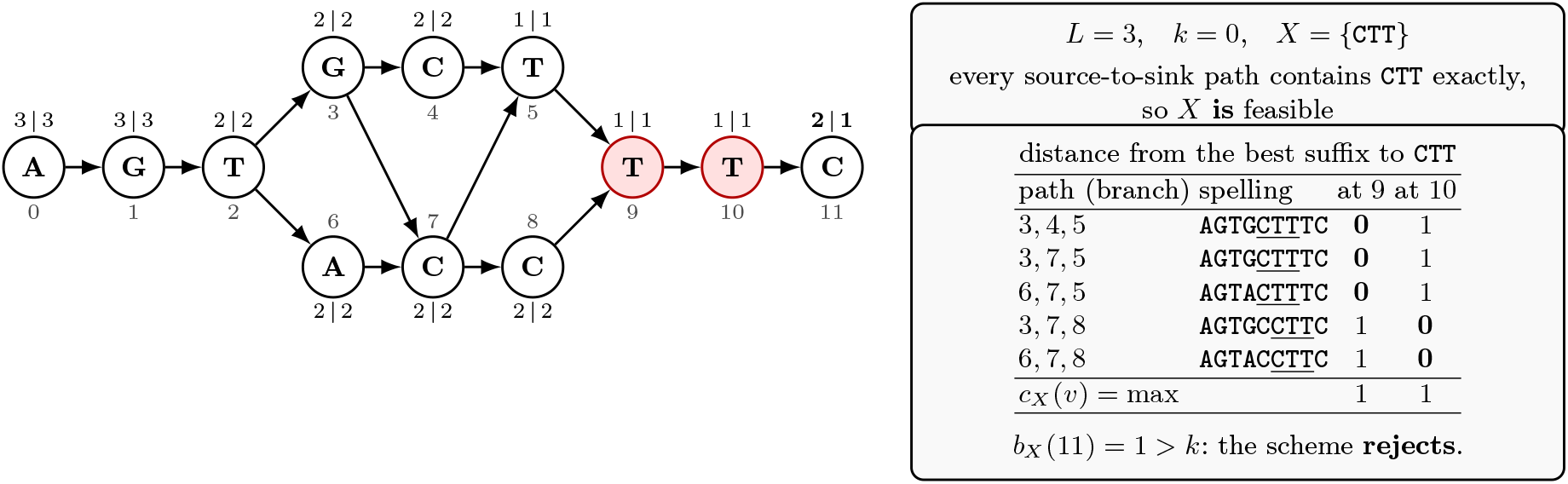
A feasible instance that the two-pass scheme rejects. Each node *v* is annotated c_*X*_(*v*) | *b*_*X*_(*v*), as defined in Fig. 2. All five source-to-sink paths contain <monospace>CTT</monospace> as an exact subpath, so *X* = <monospace>{CTT}</monospace> is feasible for *k* = 0; yet *b*_*X*_(11) = 1 > *k* and the scheme reports infeasible. The reason is visible at the shaded nodes 9 and 10, where the occurrences end: on the three paths through node 5 the match ends at 9, and on the two paths through node 8 it ends at 10, so the per-path distances are complementary. Taking the maximum over incoming paths gives *b*_*X*_ (10) = 1 at both, and no node anywhere in the graph attains *b*_*X*_ (*v*) = 0.

## 3 Experiments

### Datasets

We evaluated our algorithms on three gene-specific pangenome graphs built from their NCBI RefSeq entries [14–16] and genetic variants [17] in Clinvar [11, 12]. These specific genes, such as KRAS, were chosen for their relevance in ASO therapeutics research, respectively, cancer research [10, 22, 30, 31]. To the best of our knowledge, ready-made disease-specific target gene pangenome graphs are not available. FASTA sequences were simulated by injecting the reference sequence with gene-specific variants, s.t. there is a sequence for each variant. The directed graphs are constructed with PGGB [5] (see Appendix C on bi-directionality). In the following, the topological length is the number of nodes on a path; the sequence length is the total number of nucleotides spelled out by a path. As we require character-labeled graphs as input, topological is equal to sequence length.

Gene-specific pangenome graphs are significantly larger than oligonucleotide candidates or their target sites (12-24 bp) [4]. For ease of processing and analogous to sliding-window approach for a linear reference, we split the gene-specific pangenome graphs into acyclic subgraphs. Directed Superbubbles extracted with BubbleFinder [6, 25] are defined topologically, so their sequence lengths may be shorter than *L*; we therefore merge them into subgraphs satisfying: (i) each subgraph has a single source *s* and sink *t* whose sequences are trimmed to a suffix and a prefix of length *L* - 1, respectively, (ii) the shortest path (excluding *s*) has a sequence length of at least *L*, (iii) the sequence length of *t* is at least 3; and (iv) if *s* has a sequence length smaller *L* - 1, the graph is extended to a single prefix node s.t. condition (i) can be fulfilled. Condition (iii) was chosen empirically so that variant hotspots are merged. Condition (iv) is relevant for the lossless filter, computing the recurrence *d*_*x*_ for a string of length *L* treats at least *L* - 1 rows following the initialization row as prefix rows. Table 1 shows an overview of the pangenome graphs used in the experiments along with graph statistics of the extracted character-labeled acyclic subgraphs. For simplicity, subgraphs are referred to as graphs in the following.

**Table 1.** Summary statistics of pangenome graphs used in evaluation.

| Gene | Full graph |  |  |  |  | Acyclic subgraphs (character-labeled) |  |  |
|---|---|---|---|---|---|---|---|---|
|  | Sequences<br>[#] | Nodes<br>[#] | Edges<br>[#] | Length<br>[bp] | Edges<br>per node | Count<br>[#] | Edges per node<br>(range) | Size \|*E*\|<br>(range) |
| SOD1 | 250 | 563 | 812 | 9541 | 1.44 | 41 | 1.00–1.39 | 40–168 |
| TP53 | 3830 | 5354 | 9462 | 30647 | 1.77 | 45 | 1.00–1.64 | 40–6309 |
| KRAS | 516 | 1136 | 1666 | 46231 | 1.47 | 75 | 1.00–1.56 | 41–231 |

### Implementation

All introduced approaches are implemented in Python and are freely available at https://github.com/Schlieplab/String-DAG-Cover. The implementation for Algorithm 1 builds the product graph from a DFA and a graph disregarding nodes that include a finite state. The compressed sparse row representation for node adjacency is used to check for a remaining path. We implemented two DFAs. First, the *Aho–Corasick DFA* that builds a trie from the set of generated edit neighborhoods for each *x* ∈ X, and computes the full transition function for the finite alphabet *Σ* = {`A, C, G, T`} subsequently. Edit neighborhoods are generated by `quickmers` [9]. The resulting product graph is referred to as *AC-PG N*^*k*^(*x*) for a single candidate string *x* G *X* and *AC-PG N*^*k*^*(X*) for multiple strings X. Second, the *Ukkonen DFA* (see Definition 14) that implements the algorithm described in [26] taking a string x and a threshold k as input. The resulting product graph is referred to as *Ukk-PG N*^*k*^(*x*). The *LLF for String-DAG-Cover-Verifying through dynamic programming* implementation, referred to as *LLF DP*, expects a graph and a string x as input. As *d*_*x*_, *c*_*x*_ and *b*_*x*_ depend only on predecessors, one pass in topological order suffices.

#### Baseline comparison

The brute-force approach to solve Problem 2 is to compute *D*_*x*_ as described in Equation 2 and the bottleneck costs *B*_*x*_*(t)* by iterating over the set of paths *P. Naive edlib* uses the `edlib` library [33] to compute the edit distance in infix mode for each candidate string and path. *Naive DP* uses a naive dynamic programming algorithm.

#### Results

Algorithm 1 was evaluated for edit neighborhoods *N*^*k*^ (*x*) for 0≤*k*≤3 iteratively for each candidate string *x* ∈ *X. X* is defined as the set of all strings of length *L* = 20 with an exact match on the input graph. Therefore, the number of strings evaluated per graph vary depending on the graph size. *L* was chosen to fall well within the range common for target sites in RNA therapeutic research [4]. We categorized graphs in three categories by their node and edge count in GFA format: (1) *Small Indels* with |*E*| = 3, (2) *SNPs* with (*n* — 2) • 2 edges given *n* nodes and (3) *Other*. Our SNP criterion may also match nodes longer than one base, but no such case occurs in our data. For *Other* graphs, we focus on candidate strings with exact matches located well within the graphs. Candidate strings with exact matches on shared prefixes or suffixes are considered trivial results.

Figure 4 shows the percentage of strings *x* ∈ *X* for which Algorithm 1 returned “yes” using *Ukk-PG N*^*k*^ (*x*) for 0 < *k* < 3. SNPs are not included in the Figure, as 0% of candidate strings covered the graphs with *k* = 0 while all strings covered the graphs with *k* =1 as this was expected. The share of candidate strings covering the graphs generally declines with graph size, and the marked outliers suggest it also depends on the number of paths per graph. Note the outliers for *Small Indels* for *k* = 0 with 45 and 43 edges, in which a 6bp repeat sequences and, respectively, a T run is inserted in a T-rich region. The largest graphs returning a non-zero percentage of strings covering a graph for *N*^3^(*x*) and *N*_*2*_*(k)* have 780 and 144 edges, respectively (see Appendix Table D.2). Both graphs consist of at least 40 million paths.

**Fig 4.**
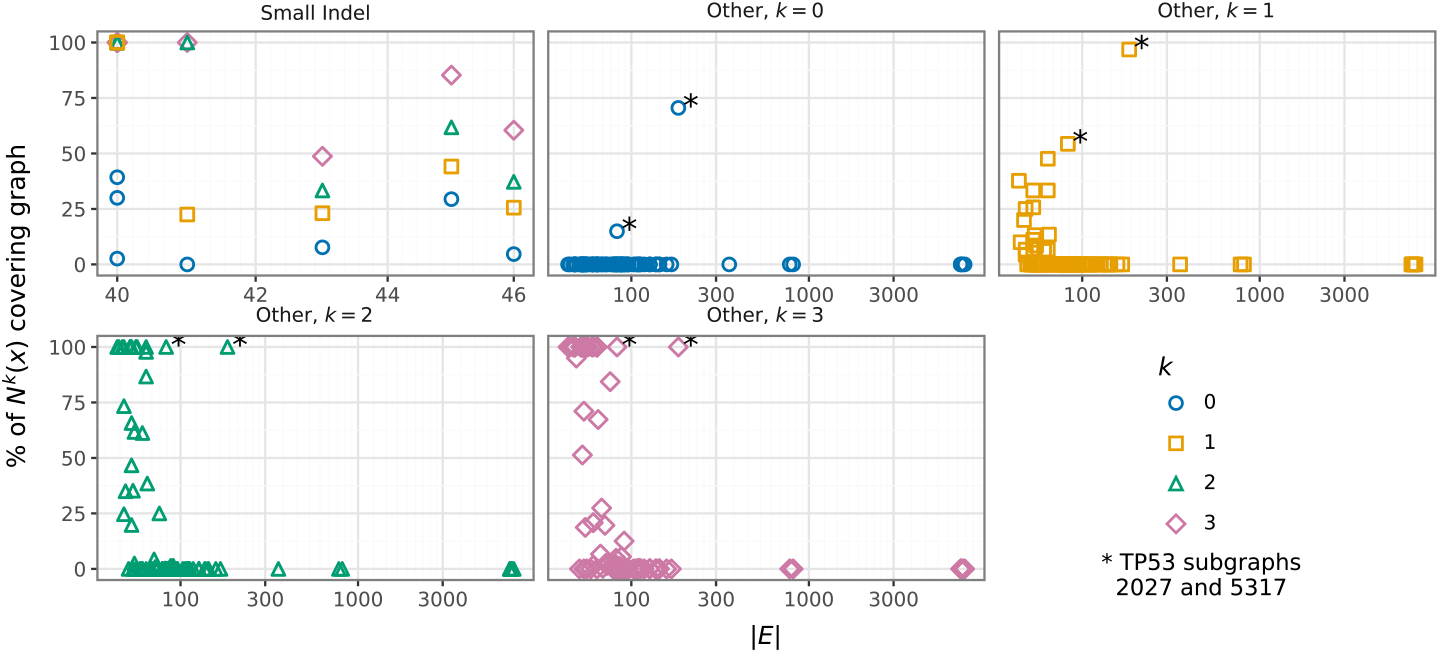
Results of *Ukk-PG N*^*k*^(*x*) for graphs of all three genes. With increasing graph size, fewer strings can cover the graphs. For many smaller graphs in *Other* there is a small percentage of candidate strings that can cover the graphs even with *k* = 1. Graphs marked with * stem from regions where a cycle-inducing branch was excluded, leaving long near-linear subgraphs with 6 and 12 paths, respectively.

#### Correctness

If Algorithm 1 returns “yes” for a string *x*, its edit neighborhood and a graph, it implies that *D*_*x*_ < *k*. Thereby, we can evaluate the correctness of our approach empirically. The smallest *k* for which Algorithm 1 returns “yes” matches *D*_*x*_ in the brute-force evaluation. If Algorithm 1 returns “no” for all 0 ≤ *k* ≤ 3, then *D*_*x*_ > 3. Over 141 graphs, that were run at least once for *Naive edlib* and *Ukk-PG N*^*k*^(*x*), and 154192 strings, all results matched. Equivalently, results for *AC-PG N*^*k*^(x) were positively assessed.

*Results on* |*X*| = 2. *AC-PG N*^*k*^(*x*) can be easily extended to include edit neighborhoods of a set of strings *X*. As the run times are significantly larger (ref. Figure 5), we selected a subset of graphs for which no string *x* ∈ *X* exists s.t. Algorithm 1 returns “yes” for *N*^2^(*x*). Here, a *lazy-build* function was used due to time constraints, constructing only the nodes and edges reachable from node (*s,u*_*in*_) (ref. Algorithm 1) of the product graph and are thereby relevant to check for a path cover. Iterating over all pairs of *X, X*^’^ = (*x,x*^’^), and their edit neighborhood *N*^2^(*X*) for graph 48 of SOD1, ∼1.6 million pairs were processed in 48 h, of which 30 pairs returned “yes” for Algorithm 1. An additional example can be found in Appendix D.3.

**Fig 5.**
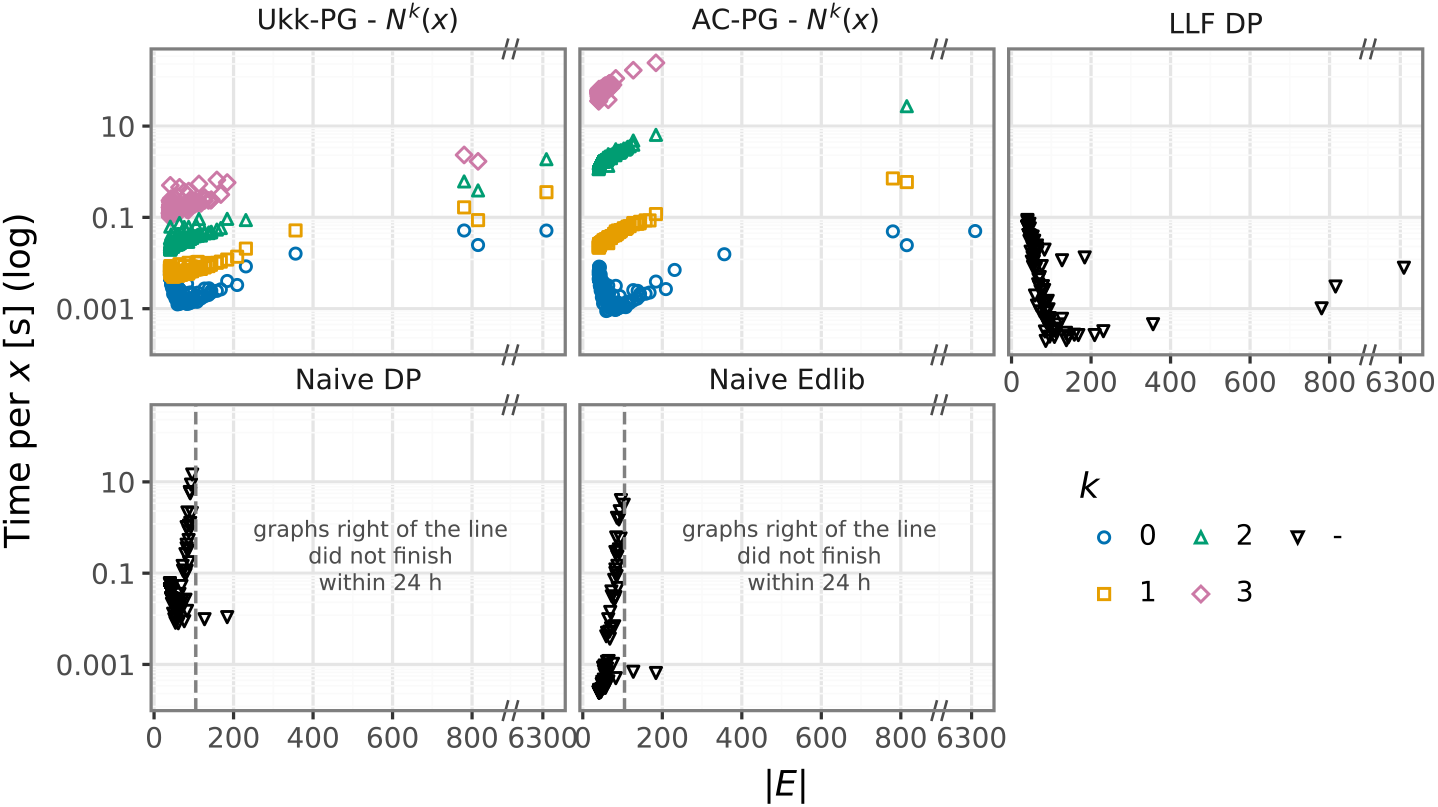
Comparison of runtime for *Ukk-PG N*^*k*^(x), *AC-PG N*^*k*^(x), *LLF DP, Naive Edlib* and *Naive DP* over the graph size |E | for all graphs over three gene-specific pangenome graphs. Total run times were limited to 24 h. The two graphs to the right of the line are large near-linear graphs with 6 and 12 paths, respectively, highlighting the dependency on the number of paths of the brute-force approaches.

#### Approximation error of LLF for String-DAG-Cover-Verifying through dynamic programming

The estimated bottleneck costs *b*_*x*_ are compared to *B*_*x*_ computed by *Naive Edlib*. Appendix Table D.3 shows approximation results per graph type including the average absolute differences and approximation ratio 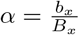 for *B*_*x*_ *>* 0. Note, that *LLF DP* does not under estimate the exact bottleneck value *B*_*x*_ and *α* is on average below 1.5 for 141 graphs evaluated.

#### Empirical run times

The introduced approaches *Ukk-PG N*^*k*^ (*x*), *AC-PG N*^*k*^(*x*), and *LLF DP* are compared to the baseline implementations *Naive Edlib* and *Naive DP*. Figure 5 show the runtime per string x in seconds. While *Naive Edlib* and *Naive DP* are faster on smaller graphs, the computations become costly as the number of paths per subgraph increases. For larger graphs, |E| ≈ 100, the product graph approaches can be faster while for even larger graphs our approaches make it feasible to evaluate the graphs within 24 h. *LLF DP* is a highly efficient alternative regardless of the graph size when approximate result are sufficient.

## 4 Discussion

We propose algorithmic approaches that can integrate the genetic diversity of genes already at early stages of the design process for oligonucleotide therapeutics. This has the potential to accelerate the design process and reduce the risk of late-stage failures. Specifically, we focused on S<sc>tring</sc>-DAG-<sc>cover-verifying</sc> approaches for a given set of potential target sites or oligonucleotides.

### Insights from experiments

Brute-force approaches show favorable performance on small graphs. However, in variant-rich regions with small edit neighborhoods their runtime exceeded 24 hours as path enumeration dominates runtime on complex graphs. In contrast, the product graph approaches showed better performance for such instances, making target site identification on gene-specific pangenome graphs feasible even when target regions are constrained to variant hotspots. The lossless filter for S<sc>tring</sc>-DAG-<sc>cover-verifying</sc> Union or the European Research Council. Neither the European using dynamic programming provides a highly efficient alternative and can serve as a quick pre-screening step for target site identification. We also presented a case, where by utilizing the Aho–Corasick DFA to encode edit neighborhoods for pairs of strings, we could verify multiple sets of strings where no single candidate alone was a viable solution for Problem 2. This suggests that secondary target sites can be successfully identified in highly diverse genetic regions when a single oligonucleotide is not sufficient. We also observed that *Ukk-PG N*^k^(*x*) outperforms *AC-PG N*^*k*^(*x*) since its DFA needs a significantly smaller number of states to encode edit neighborhoods, which consequently reduces the product graph size.

### Automata

We presented experiments for |*X*| = 2 using the Aho-Corasick DFA, but did not discuss how to solve Problem 2 when *k* > 0 and |*X*| > 1 with the Ukkonen DFA. A natural approach is an extension of Ukkonen DFA in which every state has an entry for every prefix of some string in *X* . The idea is to define state *w*^*z*^ by computing *ed*^*\**^*(y, z)* for every prefix *y* of some string in *X*. For example, if *X* consists of two strings x and x’, then *w*^*z*^ should be a vector of length |x| + |x’| + 2: the first |x| + 1 entries refer to each prefix of *x*, and the last |*x*’| + 1 entries refer to each prefix of x’. It should be possible to build the DFA by following the dynamic programming approach (based on matrix *D*) for both strings in parallel. If the strings in *X* share many prefixes, it looks sensible to reduce the length of *w*^*z*^ to | Pref(*X*)| as in Aho-Corasick DFA, even though the dynamic programming approach may become more cumbersome because the information on some prefix could be used to update multiple strings. In general, much care is needed to keep size and construction time of the DFAs tractable, both in theory and practice. A promising direction might draw inspiration from *Levenshtein DFA*. That is, DFAs *A* for which ℒ (*A*) = *N*^*k*^ *(x)* for some string *x* ∈ *Σ* ^*^ (and not *ℒ (A)* = *Σ*^*^ · *N*^*k*^(x)). For example, for a fixed k, there exists a Levenshtein DFA for x having only *O*(|*x*|) states, and such an automaton can be built in O(|*x*|) time [13, 23].

### Strengthened hardness results, approximation, and other future work

The coNP-completeness reduction on the verification problem (Theorem 4) requires a large alphabet and large *k*. We are working on tightening these in a fashion similar to the minimization problem (Theorem 3). An interesting avenue of study is the approximation of Problem 1 when *k* = 0. Adopting the greedy set cover strategy, one would obtain a logarithmic approximation guarantee on the number of paths, but it is an open problem how to implement such strategy in polynomial time. Finding a string that covers most paths may be feasible adopting the product graph techniques, but continuing the greedy scheme for finding a string that covers most of the yet uncovered paths might be tricky. As we demonstrated empirically, dynamic programming can solve a considerable number of instances. Additional finetuning of the approach with forward-backward computation should extend this to even more cases. An interesting question is whether a clean characterization of families of graphs that admit such a dynamic programming approach is possible. Also the formal proof of Problem 19 stated in Appendix E is still open.

### Applications

Here we focused on problems arising from computational design of therapeutic oligonucleotides. The inclusion into design pipelines such as ASOkai [3] and evaluation results in such settings will be reported elsewhere. Note, there are additional experimental uses for oligonucleotides from pathogen detection to primers for amplicon sequencing which likely can benefit from variation-aware design.

## Acknowledgments

The first author gratefully acknowledges funding from BTU’s graduate research school for a research stay at the University of Helsinki. The second author is funded by the Helsinki Institute for Information Technology (HIIT). Co-funded by the Research Council of Finland grants 358744, 374907. Co-funded by the European Union (ERC, SCALEBIO, 101169716). Views and opinions expressed are however those of the author(s) only and do not necessarily reflect those of the European Union or the European Research Council. Neither the European

## Footnotes

3 The proof of Theorem 4 (included in Appendix) was found with Claude Opus 5.5. All the technical content in the main article is by the authors. For a more detailed statement of AI usage, see p. 10 in the Acknowledgments.

