## Appendix for "Algorithms for Designing Therapeutic Oligonucleotides on Pangenome Graphs"

#### A Hardness proofs

First, we give the main NP-hardness proof of String-DAG-cover, by a reduction from Set Cover, in a relaxed setting where the DAG may have several sources and sinks. Then, we show that this reduction can be strengthened to prove hardness also for DAGs whose nodes are labeled from a ternary alphabet only, and such that the nodes all lie on a single path; in particular, such DAGs have a unique source and a unique sink, as required in Problem 1.

We prove hardness of the *decision version* of String-DAG-cover, which is additionally given an integer $B$ and asks whether there exists a set $X$ of strings of length $L$ with $|X| \leq B$ such that each source-to-sink path of $G$ has a subpath spelling a string that has edit distance at most $k$ from at least one string in $X$; we then say that $X$ is *feasible*, and that $X$ *covers* each such path. NP-hardness of the decision version implies NP-hardness of Problem 1, since the minimum size of a feasible $X$ is at most $B$ iff the decision version answers yes.

**Lemma 18.** *String-DAG-cover is NP-hard already for $k = 0$, if the DAG $G$ is allowed to have several sources and sinks (and all of its source-to-sink paths must be covered).*

*Proof.* We reduce from the decision version of Set Cover. Let $\mathcal{I} = (U, \mathcal{S}, B)$ be an instance of Set Cover, where $U = \{u_1, \ldots, u_n\}$ is the universe, $\mathcal{S} = \{S_1, \ldots, S_m\}$ is a family of subsets of $U$, and $B$ is the budget. The question is whether there exists an index set $J \subseteq \{1, \ldots, m\}$ with $|J| \leq B$ such that

$$U = \bigcup_{j \in J} S_j.$$

We construct an instance of String-DAG-cover. For every set $S_j \in \mathcal{S}$, introduce a distinct character $\sigma_j$. We set $L = 1$ and $k = 0$. Thus, a candidate string is just a single character, and it covers a path exactly when that character occurs on the path.

For every element $u_i \in U$, we create one directed path $P_i$. The path $P_i$ contains one node labeled $\sigma_j$ for every set $S_j$ that contains $u_i$, that is, for every $j$ such that $u_i \in S_j$. The nodes on $P_i$ can be arranged in any fixed order. The graph $G$ is the disjoint union of these paths. Since a disjoint union of directed paths is a DAG, $G$ is a DAG. The construction is clearly polynomial in the size of the Set Cover instance. Figure A.1 shows an example.

We claim that the Set Cover instance has a cover of size at most $B$ iff the constructed String-DAG-cover instance has a feasible candidate set $X$ of size at most $B$.

For the forward implication, suppose that there exists a set cover $J \subseteq \{1, \ldots, m\}$ with $|J| \leq B$. Define

$$X = \{\sigma_j : j \in J\}.$$

Consider any element path $P_i$. Since $J$ is a set cover, there exists some $j \in J$ such that $u_i \in S_j$. By construction, $P_i$ contains a node labeled $\sigma_j$. Since $L = 1$ and $k = 0$, the length-1 subpath spelling $\sigma_j$ matches the candidate string $\sigma_j$ exactly. Hence $P_i$ is covered. Therefore every path is covered by $X$, and $|X| = |J| \leq B$.

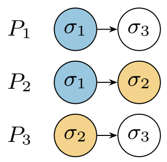


**Fig. A.1:** The reduction in the proof of Lemma 18 for $U = \{u_1, u_2, u_3\}$, $S_1 = \{u_1, u_2\}$, $S_2 = \{u_2, u_3\}$, $S_3 = \{u_1, u_3\}$. The set cover $J = \{1, 2\}$ gives $X = \{\sigma_1, \sigma_2\}$, whose colored occurrences cover all paths.

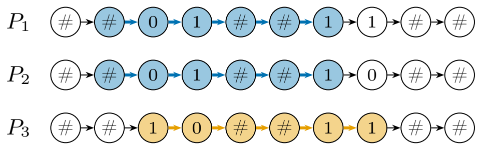


**(a)** The graph $G$, a disjoint union of paths.

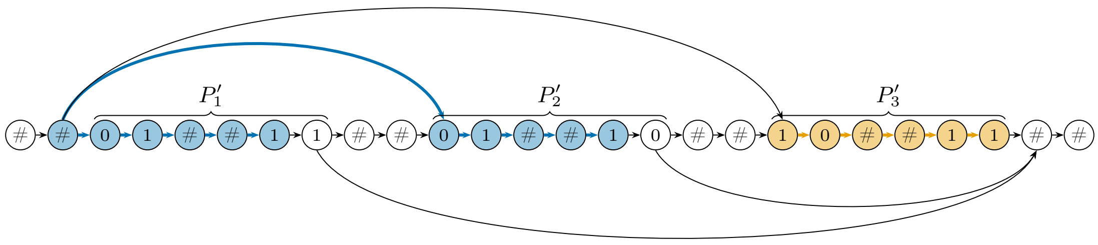


**(b)** The graph $G'$, whose nodes all lie on a single path.

**Fig. A.2:** The reduction in the proof of Theorem 3 for the instance of Figure A.1, so $T = 2$, $b(1) = 01$, $b(2) = 10$, $b(3) = 11$, and $L = 6$. In (a), the colored occurrences show that $X = \{\#01\#\#1,\ 10\#\#11\}$ covers all paths, even though its candidates are not aligned with the delimiters; the reverse direction maps them to $j(c) = 1$ and $j(c) = 2$, respectively, giving the set cover $J = \{1, 2\}$. (The forward direction would build $X = \{\#\#01\#\#,\ \#\#10\#\#\}$ from $J$.) In (b), the same three occurrences are shown in $G'$, on the source-to-sink paths spelling $P_1$, $P_2$, and $P_3$; the occurrence on $P_2$ uses the edge from the first delimiter to $P'_2$.

For the reverse implication, suppose that the constructed String-DAG-cover instance has a feasible candidate set $X$ with $|X| \leq B$. Because $L = 1$, every candidate is a single character. Moreover, any character not among $\{\sigma_1, \dots, \sigma_m\}$ does not occur in the graph and therefore covers no path, so we may assume without loss of generality that $X \subseteq \{\sigma_1, \dots, \sigma_m\}$. Define

$$J = \{j : \sigma_j \in X\}.$$

Then, $|J| = |X| \leq B$. Now consider any element $u_i \in U$. Since the path $P_i$ is covered, it contains some label $\sigma_j \in X$. By construction, the label $\sigma_j$ appears on $P_i$ only if $u_i \in S_j$. Therefore, for every $u_i \in U$, there exists some $j \in J$ such that $u_i \in S_j$. Hence

$$U = \bigcup_{j \in J} S_j,$$

so $J$ is a set cover of size at most $B$.

Thus, the Set Cover instance is a yes-instance iff the constructed String-DAG-cover instance has a feasible solution of size at most $B$. Therefore the decision version of String-DAG-cover is NP-hard. Consequently, the optimization version of String-DAG-cover is NP-hard as well.

**Theorem 3.** *String-DAG-cover is NP-hard already for $k = 0$, even on DAGs whose nodes are labeled from an alphabet of size three, and whose nodes all lie on a single path (i.e. they consist of a single path, plus additional edges).*

*Proof.* We modify the proof of Lemma 18 as follows. We will use the ternary alphabet $\{0, 1, \#\}$.

First, instead of using a single node labeled $\sigma_j$ for every set $S_j$, we use a sequence of $T$ nodes, where $T = \lfloor \log_2 m \rfloor + 1$ is the number of bits needed to represent all set indices between 1 and $m$. These $T$ nodes spell the binary encoding $b(j)$ of each index $j \in \{1, \dots, m\}$ (padded with 0s to have exactly $T$ bits); we call them a *block*.

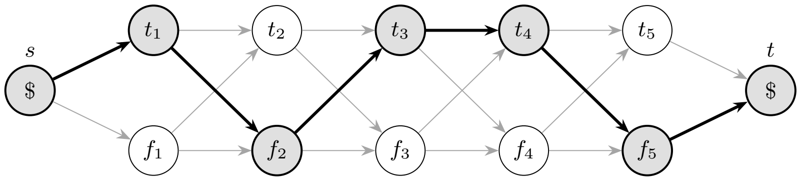


| | | | | | | |
|---|---|---|---|---|---|---|
| $S =$ | $t_1$ | $f_2$ | $t_3$ | $t_4$ | $f_5$ | |
| $c_j =$ | $\psi$ | $f_2$ | $t_3$ | $\psi$ | $f_5$ | |
| cost | 1 | 0 | 0 | 1 | 0 | total $= n - 3 = k$ |

**Fig. A.3:** The reduction of Theorem 4 for $n = 5$ and the clause $D_j = (x_2 \vee \neg x_3 \vee x_5)$, whose falsifying characters are $f_2, t_3, f_5$. Each diamond chooses the value of one variable, so source-to-sink paths correspond bijectively to assignments; the highlighted path is $\alpha = (\mathrm{T}, \mathrm{F}, \mathrm{T}, \mathrm{T}, \mathrm{F})$, and $S$ is its subpath without the source $s$ and the sink $t$, which are both labeled \$. The pattern $c_j$ carries the three falsifying characters in their own positions and the character $\psi \notin \lambda(V)$ elsewhere. Each $\psi$ costs one edit no matter what, so the edit distance is at least $n-3$ always, with equality exactly when the three named characters match—that is, exactly when $\alpha$ falsifies $D_j$. Setting $k = n - 3$ therefore makes a path uncovered iff its assignment satisfies every clause.

Second, to control matches of candidate strings that could overlap consecutive blocks, we use a delimiter $D$ which is a path of $T$ nodes labeled with #. We construct each path $P_i$ as $D$, then the block of the first set containing $u_i$, then again $D$, then the block of the second set containing $u_i$, and so on, until ending again with $D$. As in the proof of Lemma 18, the graph $G$ is the disjoint union of the paths $P_1, \ldots, P_n$. We use again $k = 0$, but now we set $L = 3T$. See Figure A.2(a) for an example.

We claim that the Set Cover instance has a cover of size at most $B$ iff the constructed instance has a feasible candidate set of size at most $B$.

For the forward implication, given a set cover $J$, let $X = \{\#^T\, b(j)\, \#^T : j \in J\}$; these strings have length $L$. For every $P_i$, there is $j \in J$ with $u_i \in S_j$, and then $\#^T\, b(j)\, \#^T$ is a substring of $P_i$, so $P_i$ is covered.

For the reverse implication, let $X$ be feasible with $|X| \leq B$. Consider any $c \in X$ that occurs in some $P_i$. Since $L = 3T$, $c$ contains $b(j)$ for some $1 \leq j \leq m$. Let $j(c)$ be the smallest such $j$, and let $J = \{j(c) : c \in X\}$, so $|J| \leq B$. Now, for every $u_i$, the path $P_i$ is covered by some $c \in X$. Since $b(j(c))$ occurs in $c$, it occurs in $P_i$, and so we get $u_i \in S_{j(c)}$. Hence $J$ is a set cover.

Finally, we connect the paths so that all nodes lie on a single path. Write $P_i = D\, P'_i\, D$, that is $P'_i$ is the middle part of $P_i$ except the delimiters at its two ends. Let $G'$ be the path

$$D,\ P'_1,\ D,\ P'_2,\ D,\ \ldots,\ D,\ P'_n,\ D,$$

plus the edges from the last node of the first delimiter to the first node of every $P'_i$, and from the last node of every $P'_i$ to the first node of the last delimiter. All its source-to-sink paths spell exactly all the strings $D\, P'_i\, D\, P'_{i+1}\, D \cdots D\, P'_{i'}\, D$, for $1 \leq i \leq i' \leq n$. Figure A.2(b) shows $G'$ for the example above. The above claim still holds for $G'$. If $X$ covers every $P_i$, then it covers every source-to-sink path of $G'$, since each such path contains $D\, P'_i\, D = P_i$ for some $i$. Conversely, if $X$ is feasible for $G'$, then in particular it covers the source-to-sink paths spelling $D\, P'_i\, D = P_i$, for every $i$, and the reverse argument above applies unchanged.

Unlike $G$, the DAG $G'$ has a unique source (the first node of the first delimiter) and a unique sink (the last node of the last delimiter), so it is a valid input to Problem 1. Therefore the decision version of String-DAG-cover is NP-hard on such DAGs, and so is Problem 1.

**Theorem 4.** *String-DAG-cover-verifying is coNP-complete.*

We write Uncovered for the complement of String-DAG-cover-verifying: does there exist a source-to-sink path none of whose subpaths is covered?

*Proof. Membership.* An uncovered source-to-sink path certifies UNCOVERED: it has $O(|V|^2)$ subpaths, each checkable against each $c \in X$ in $O(L|V|)$ time.

*Hardness.* We reduce from 3-SAT with variables $x_1, \ldots, x_n$ ($n \geq 3$) and clauses $D_1, \ldots, D_r$, each over three distinct variables. Let $\Sigma = \{t_1, f_1, \ldots, t_n, f_n\} \cup \{\psi, \$\}$. The graph is a chain of $n$ diamonds: for each $i$ create nodes $T_i$ with $\lambda(T_i) = t_i$ and $F_i$ with $\lambda(F_i) = f_i$, and add all four edges from $\{T_i, F_i\}$ to $\{T_{i+1}, F_{i+1}\}$ for $i < n$. Finally, add a source $s$ with edges to $T_1, F_1$ and a sink $t$ with edges from $T_n, F_n$, with $\lambda(s) = \lambda(t) = \$$. Thus, source-to-sink paths correspond bijectively to truth assignments: $P_\alpha$ spells $\$\, b_1 \cdots b_n \,\$$ with $b_i = t_i$ iff $\alpha(x_i) = \text{true}$.

Set $L = n$ and $k = n - 3$. For a clause $D_j$ over variables $i_1 < i_2 < i_3$ let $c_j = \beta_1 \cdots \beta_n$, where $\beta_{i_p}$ is the character *falsifying* the $p$-th literal of $D_j$ (that is $f_{i_p}$ for a positive literal and $t_{i_p}$ for a negative one) and $\beta_i = \psi$ otherwise. Put $X = \{c_1, \ldots, c_r\}$. Figure A.3 illustrates the reduction.

The character $\psi$ does not occur in $G$, so in any alignment of a subpath $S$ with $c_j$ each of the $n-3$ occurrences of $\psi$ is deleted or substituted, costing at least one each; hence the edit distance of $S$ and $c_j$ is at least $n - 3 = k$ always.

Suppose the edit distance of $S$ and $c_j$ is $k$. Then the budget is consumed entirely by the $\psi$ positions and every other operation is free: the three characters $\beta_{i_1}, \beta_{i_2}, \beta_{i_3}$ are matched exactly and no character of $S$ is left unaligned. In particular $S$ contains $\beta_{i_1}, \beta_{i_2}, \beta_{i_3}$ in this order. As the characters $t_i, f_i$ are variable-specific, $\beta_{i_p}$ occurs in $P_\alpha$ only as $b_{i_p}$, whence $b_{i_p} = \beta_{i_p}$ for $p = 1, 2, 3$, that is, $\alpha$ falsifies $D_j$.

Conversely, if $\alpha$ falsifies $D_j$ then $b_{i_p} = \beta_{i_p}$ for all $p$, and aligning the subpath $S = b_1 \cdots b_n$ of $P_\alpha$ (i.e., without $s$ and $t$) with $c_j$ position by position substitutes exactly the $n - 3$ characters facing $\psi$, so the edit distance of $S$ and $c_j$ is $k$.

Therefore $P_\alpha$ is uncovered iff $\alpha$ satisfies every clause, and UNCOVERED is NP-hard.

### B Proofs of Aho–Corasick graph product

#### B.1 Proof of Theorem 7

*Proof.* To prove the theorem 7, we will show that Algorithm 1 returns "no" iff there exists an $s$-$t$ path of $G$ that contains no subpath spelling a string in $Y$.

($\Rightarrow$) Assume that Algorithm 1 returns "no". Then, $(s, q_{in}) \in V^*_{\mathcal{A}}$ and there exists $u \in Q$ such that (i) $(t, u) \in V^*_{\mathcal{A}}$ and (ii) $G^*_{\mathcal{A}}$ contains a walk from $(s, q_{in})$ to $(t, u)$. This means that, for some $h \geq 2$, there exist nodes $(v_1, q_1)$, $(v_2, q_2)$, ..., $(v_h, q_h)$ in $V^*_{\mathcal{A}}$ such that (i) $(v_1, q_1) = (s, q_{in})$, (ii) $((v_{i-1}, q_{i-1}), (v_i, q_i))$ is an edge of $G^*_{\mathcal{A}}$ for every $2 \leq i \leq h$ and (iii) $(v_h, u_h) = (t, u)$. By the definition of $G^*_{\mathcal{A}}$, (i) for every $1 \leq i \leq h$ we have $q_i \notin F$ and (ii) for every $2 \leq i \leq h$ we have $((v_{i-1}, q_{i-1}), (v_i, q_i)) \in E_{\mathcal{A}}$. By the definition of $E_{\mathcal{A}}$, for every $2 \leq i \leq h$ we have $(v_{i-1}, v_i) \in E$ and $q_i = \delta(q_{i-1}, \lambda(v_{i-1}))$. In particular, $(v_i)_{1 \leq i \leq h}$ is an $s$-$t$ path in $G$, and we have $\delta(q_{in}, \lambda(v_1)\lambda(v_2) \ldots \lambda(v_{i-1})) = q_i$ for every $1 \leq i \leq h$.

The conclusion will follow if we show that no subpath of $(v_i)_{1 \leq i \leq h}$ spells a string in $Y$. Suppose, for the sake of contradiction, that there exists $y \in Y$ spelling a subpath of $(v_i)_{1 \leq i \leq h}$. This means that there exists $1 \leq i \leq h - |y| + 1$ such that $\lambda(v_i)\lambda(v_{i+1})\lambda(v_{i+2}) \ldots \lambda(v_{i+|y|-1}) = y$. We cannot have $i = h - |y| + 1$; otherwise, we would have $v_{i+|y|-1} = v_h = t$, so $\lambda(v_{i+|y|-1})) = \lambda(t) = \$$ and the last character of $y$ would be \$, which is a contradiction. Consequently, we have $1 \leq i \leq h - |y|$, and so $i + |y| - 1 \leq h - 1$. In particular, $y$ is a suffix of $\lambda(v_1)\lambda(v_2) \ldots \lambda(v_{i+|y|-1})$, so $\lambda(v_1)\lambda(v_2) \ldots \lambda(v_{i+|y|-1}) \in \Sigma^* \cdot Y$. Since $\mathcal{L}(\mathcal{A}) = \Sigma^* \cdot Y$, we have $\delta(q_{in}, \lambda(v_1)\lambda(v_2) \ldots \lambda(v_{i+|y|-1})) \in F$, or equivalently, $q_{i+|y|} \in F$. This is a contradiction because $q_{i+|y|} \notin F$.

($\Leftarrow$) Assume that there exists an $s$-$t$ path $(v_i)_{1 \leq i \leq h}$ in $G$ that contains no subpath spelling a string in $Y$. In particular, $v_1 = s$ and $v_h = t$.

Let us prove that $(s, q_{in}) \in V^*_{\mathcal{A}}$. Suppose, for the sake of contradiction, that $(s, q_{in}) \notin V^*_{\mathcal{A}}$. By the definition of $V_{\mathcal{A}}$ we have $(s, q_{in}) \in V_{\mathcal{A}}$, so by the definition of $V^*_{\mathcal{A}}$ we have $q_{in} \in F$, which implies that the empty string $\epsilon$ is in $\mathcal{L}(\mathcal{A})$. From $\mathcal{L}(\mathcal{A}) = \Sigma^* \cdot Y$ we obtain $\epsilon \in Y$. This is a contradiction because $\epsilon$ spells the empty subpath of $(v_i)_{1 \leq i \leq h}$.

Define $q_1 = q_{in}$. For every $2 \leq i \leq h$, recursively define $q_i = \delta(q_{i-1}, \lambda(v_{i-1}))$. By the definition of $E_{\mathcal{A}}$ we have $((v_{i-1}, q_{i-1}), (v_i, q_i)) \in E_{\mathcal{A}}$ for every $2 \leq i \leq h$. In particular $(v_i, q_i)_{1 \leq i \leq h}$ is a path in $G_{\mathcal{A}}$ starting at $(s, q_{in})$ and ending at $(t, q_h)$. Moreover, $\delta(q_{in}, \lambda(v_1)\lambda(v_2) \ldots \lambda(v_{i-1})) = q_i$ for every $1 \leq i \leq h$.

We only need to prove that $q_i \notin F$ for every $1 \leq i \leq h$ because then we can conclude that (i) $(t, q_h) \in V^*_{\mathcal{A}}$ and (ii) $(v_i, q_i)_{1 \leq i \leq h}$ is a walk in $G^*_{\mathcal{A}}$ starting at $(s, u_{in})$ and ending at $(t, u_q)$, so Algorithm

1 returns "no". Suppose, for the sake of contradiction, that for some $1 \leq j \leq h$ we have $q_j \in F$. Since $\delta(q_{in}, \lambda(v_1)\lambda(v_2)\ldots\lambda(v_{j-1})) = q_j$, we have $\lambda(v_1)\lambda(v_2)\ldots\lambda(v_{j-1}) \in \mathcal{L}(\mathcal{A})$. From $\mathcal{L}(\mathcal{A}) = \Sigma^* \cdot Y$ we obtain that there exists $y \in Y$ such that $\lambda(v_{j-|y|})\lambda(v_{j-|y|+1})\ldots\lambda(v_{j-1}) = y$. Hence, the subpath $(v_i)_{j-|y|\leq i\leq j-1}$ of $(v_i)_{1\leq i\leq h}$ spells $y$, which is a contradiction. ◻

##### B.2 Proof of Lemma 8

*Proof.* We first build $G_\mathcal{A}$ by (i) building $V_\mathcal{A}$ in $O(|V| \cdot |Q|) \subseteq O(|E| \cdot |Q|)$ time and then (ii) building $E_\mathcal{A}$ in $O(|E| \cdot |Q|)$ time through the bijection $\phi : E \times Q \to E_\mathcal{A}$ of Remark 6 (in particular, $|E_\mathcal{A}| = |E| \cdot |Q|$). Next, by scanning $G_\mathcal{A}$, in $O(||E_\mathcal{A}|) \subseteq O(|E| \cdot |Q|)$ time we build $G^*_\mathcal{A}$. Lastly, within the same time bound we can check whether $(s, q_{in}) \in V^*_\mathcal{A}$ and traverse $G^*_\mathcal{A}$ to check the condition in Line 3.

##### B.3 Proof of Corollary 13

*Proof.* We first build $\mathcal{A}_X$ in $O(\sum_{x\in X} |x|)$ time (Lemma 12). We know that $\mathcal{L}(\mathcal{A}_X) = \Sigma^* \cdot X$ (Corollary 11), so we can apply Theorem 7 choosing $Y = X$. This means that Algorithm 1 decides whether each $s$-$t$ path of $G$ has a subpath spelling a string in $X$. By definition, $\mathcal{A}_X$ has $|\text{Pref}(X)| \leq \sum_{x\in X} |x|$ states, so by Lemma 8, Algorithm 1 can be implemented in $O(|E| \cdot \sum_{x\in X} |x|)$ time.

##### B.4 Proof of Corollary 17

*Proof.* We first build $\mathcal{U}^k_x$ in $O(|x|^{k+2})$ time (Lemma 16). We know that $\mathcal{L}(\mathcal{U}^k_x) = \Sigma^* \cdot N^k(x)$ (Lemma 15), so we can apply Theorem 7 choosing $Y = N^k(x)$. This means that Algorithm 1 decides whether each $s$-$t$ path of $G$ has a subpath spelling a string that has edit distance at most $k$ from $x$. By Lemma 16, $\mathcal{U}^k_x$ has $O(|x|^{k+1})$ states, so by Lemma 8, Algorithm 1 can be implemented in $O(|E| \cdot |x|^{k+1})$ time.

#### C Bubble extraction on bidirectional graphs

PGGB builds bidirected pangenome graphs. Even though our gene-specific graphs are directed by nature, the bubble/subgraph extraction can be easily modified for bidirectional graphs. It has been shown that the majority of snarls in human pangenome graphs are ultrabubbles [20]. Instead of extracting directed superbubbles, ultrabubbles are identified that can be subsequently converted into (directed) superbubbles [6].

#### D Additional results

All detailed results and the scripts to reproduce the plots can be accessed here on Zenodo (DOI: `10.5281/zenodo.23926887`).

##### D.1 Details for *Ukk-PG* $N^k(x)$

Tables D.1 and D.2 list details of the graphs showing significant results described in the main paper.

**Table D.1:** Small Indel graphs showing a higher percentage of candidate strings covering the graphs in the exact case $k = 0$

| Gene | GFA Coordinates | Graph size $\|E\|$ | Sequence (parentheses mark the indel) |
|---|---|---|---|
| TP53 | $53 \to 55$ | 45 | `...TGTGCCA(GCCGTG)GCCGTGG...` |
| SOD1 | $555 \to 557$ | 43 | `...TTGTCAGAA(TTTC)TTTGTCA...` |

**Table D.2:** Large graphs for which candidate strings were found to cover the graph with small edit neighborhoods $k$.

| Gene | GFA coordinates | Graph size $\lvert E\rvert$ | Candidate strings [#] | Strings covering graph [#] | $k$ |
|---|---|---|---|---|---|
| TP53 | 2557 → 2563 | 780 | 24773 | 4 | 3 |
| KRAS | 852 → 924 | 144 | 73347 | 16 | 2 |

#### D.2 Result plot for *AC-PG* $N^k(x)$

For completion, the results for *AC-PG* $N^k(x)$ are shown in Figure D.1. While showing fewer data points (not all graphs could be evaluated due to increased runtimes), the trends are analogous to *Ukk-PG* $N^k(x)$ as also implied by the evaluated correctness.

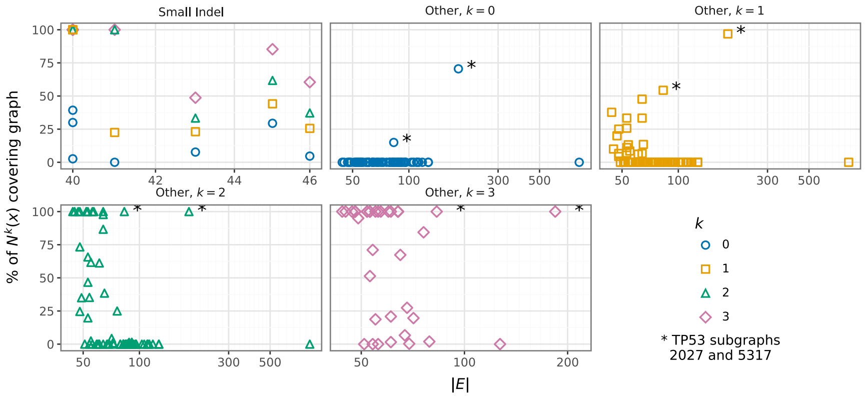


**Fig. D.1:** Results of *AC-PG* $N^k(x)$ for graphs of all three genes. Note, as not all graphs could be run in 24 h, the x-axis scale for graph size $|E|$ is limited. Generally, the same trend as for *Ukk-PG* $N^k(x)$ can be observed. Graphs marked with $*$ are larger near-linear graphs with 6 and 12 paths, respectively.

#### D.3 Additional results for $|X| = 2$

Additionally to graph 48 of SOD1, we found graph 38 of the TP53 pangenome graph that showed positive results for a set of strings $|X| = 2$. 428 275 pairs of candidate strings were evaluated in $\approx 27.3$ h, where 144 pairs returned "yes" for Algorithm 1.

#### D.4 Approximation error of lossless filter for String-DAG-Cover-Verifying through dynamic programming

Table D.3 shows the approximation results for *LLF DP* by graph type. While SNPs have no approximation error, the approximation is reasonably small otherwise.

#### D.5 Benchmarks

Measurements with a short total runtime were repeated multiple times, and we report the mean over all runs; graphs with a longer total runtime were run once. Runs that did not finish within 24 h were stopped and are not shown. Benchmarks were performed on Slurm cluster nodes with 64 cores each, single process. Each graph with exclusively allocated node, so concurrent benchmarks did not share compute resources.

**Table D.3:** Approximation results for *LLF DP* compared to the *Naive edlib* baseline. The results include 141 graphs. While the average approximation factor $\alpha = \frac{b_x}{B_x}$, for $B_x > 0$, is below 1.5, the maximum $\alpha$ can be higher. Maximum $\alpha$ for *Small Indels* correlates with the length of insertion.

| Graph type | Graphs [#] | Strings [#] | $d_x \neq D_x$ [%] | Avg. $\lvert d_x - D_x \rvert$ | Avg. $\frac{\lvert d_x - D_x \rvert}{L}$ | $d_x < D_x$ [%] | Zero mismatch [#] | Avg. $\alpha$ | Max. $\alpha$ |
|---|---|---|---|---|---|---|---|---|---|
| Small Indels | 7 | 252 | 30.6 | 0.64 | 0.03 | 0 | 36 | 1.25 | 6 |
| SNPs | 72 | 2940 | 0.0 | 0.0 | 0.0 | 0 | 0 | 1.0 | 1 |
| Other | 62 | 151000 | 40.7 | 0.51 | 0.03 | 0 | 19 | 1.07 | 4 |

### E Extension of Problem 2 to recognize subpaths spelling strings of fixed length $L$

So far we allowed the sub path length to be variable. In particular for the short length of therapeutic oligonucleotides, e.g. Anti-Sense Oligonucleotides (ASOs) and small interfering RNAs (siRNAs), the length of the oligonucleotides is limited due to constraints from therapeutic application and typically falls in the range of 12-24 bp [4]. Consequently, the length of binding sites of oligonucleotides in the mRNA of the target gene is also limited; e.g. oligonucleotides binding to the base of mRNA stem-loops seems thermodynamically unrealistic. The consideration of fixed $L$ is thus a reflection of the intended application domain leading to the introduction of the following problem:

*Problem 19 (String-DAG-cover-verifying-fixed-length).* Given a DAG $G$, a target length $L$, a set $X$ of strings of length $L$, and an edit distance threshold $k$, decide whether each $s$-$t$ path of $G$ has a subpath spelling a string that has length $L$ and has edit distance at most $k$ to at least one string in $X$.

The following results are empirical. No formal correctness guarantee is provided for this variant; this is left for future work.

#### E.1 Implementation

**Aho–Corasick.** For the solution to Problem 19, the implementation of Aho–Corasick implementation does not change. The edit neighborhood restricting its members to a fixed length $L$, $N^k(x|L)$, can be generated with `quickmers` [9], which provides a function to generate edit neighborhoods of fixed length, namely the length of the string to generate the edit neighborhood for. The Aho–Corasick automaton is constructed subsequently.

**Ukkonen automaton.** Note, Ukkonen's automaton for $N^k(x)$ recognizes every string $x'$, s.t. $ed(x, x') \leq k$ and $|x'| \neq L$. To assure we only recognize subpaths of length $L$ for $|X| = 1$ we need to change two aspects of the implementation for Problem 19. (1) Given a matrix $D = (d_{ij})$, testing whether two strings have an edit distance of $ed(a, b) \leq k$, the values in $D$ are monotonically increasing along the diagonals of $D$ [26]. Furthermore, any $d_{ij} > k$ cannot be part of the path leading to the final edit distance and any $d_{ij} \leq k$ form a band around the main diagonal. By keeping track which position in the column vector of a state in the DFA is on the diagonal we can enforce the condition to only match subpaths of length $L$. In a global alignment of two strings of equal length, the main diagonal consists of all cells $d_{ij}$ with $i = j$. The cost for an insertion, and thereby moving away from the diagonal where $i \neq j$, implies a deletion at a later stage of the edit distance computation, and vice versa. We implemented a pruning step, that by tracking the position on the diagonal, or number of characters of a string read, evaluates which position on the column vector implies a future insertion/deletion. If this estimated edit cost, the current edit operations and the expected edit operations, exceeds $k$ this position can be pruned. (2) A state is now specific to $l$, the number of characters read, and therefore, the extended Ukkonen automaton can only recognize subpaths that are prefixes of each path of a graph. By redefining states to set of $l$-specific column vectors, the initial column $d_{i0}$, with $l = 0$, can be iteratively added to each state set that is explored, thereby allowing the pattern to start from any

**Table E.1:** Approximation results for *LLF DP* $x|L$ compared to results from the *Naive DP* $x|L$ baseline. The results include a total of 140 graphs. While the average approximation factor $\alpha = \frac{b_x}{B_x}$, for $B_x > 0$, is below 1.6, the maximum $\alpha$ can be higher.

| Graph type | Graphs [#] | Strings [#] | $d_x \neq D_x$ [%] | Avg. $\lvert d_x - D_x \rvert$ | Avg. $\frac{\lvert d_x - D_x \rvert}{L}$ | $d_x < D_x$ [%] | Zero mismatch [#] | Avg. $\alpha$ | Max. $\alpha$ |
|---|---|---|---|---|---|---|---|---|---|
| Small Indels | 7 | 252 | 42.9 | 1.37 | 0.07 | 0 | 36 | 1.53 | 8 |
| SNPs | 72 | 2940 | 0 | 0 | 0 | 0 | 0 | 1 | 1 |
| Other | 61 | 111712 | 40.4 | 0.7 | 0.04 | 0 | 19 | 1.09 | 6 |

position in the graph. A similar concept using state sets was introduced by Rabin and Scott [21] where the concept of subset construction was used to convert non-deterministic state automata to deterministic state automata.

**LLF** To extend the dynamic programming-based lossless filter to a fixed alignment window $L$, we introduce an additional dimension, s.t. $d_{x,L}(v, i, l)$ is computed, where $0 \leq l \leq i \leq L$. Now let $c_{x,L}(v) = d_{x,L}(v, L, L)$. The bottleneck costs $b_x(t)$ are estimated equivalently as described in Section 2.5.

#### E.2 Experiments

**Baseline comparison** For evaluating Problem 19, the naive implementation is extended to compute the best edit distance to each path of the graph constraining an alignment window to length $L$, labeled in Fig. E.3 as *Naive DP* $x|L$.

**Results.** Equivalently, we evaluate the results and empirical run times for solving Problem 19. The Ukkonen graph product *Ukk-PG* $N^k(x|L)$ and the Aho–Corasick product graph, *AC-PG* $N^k(x|L)$, are compared with the *Naive DP* $x|L$ and lossless filter through dynamical programming *LLF* $x|L$. Figure E.3 shows a similar trend compared to the evaluation of Problem 2. At the time of submission, not all graphs could be evaluated ensuring a total runtime of 24 h. Note, while the runtimes for the *Ukk-PG* $N^k(x|L)$ increase compared to *Ukk-PG* $N^k(x)$ due to the increased number of states in the Ukkonen DFA which is in turn a result of the modified implementation described above. Therefore the size of the product graph increases. The run times for *AC-PG* $N^k(x|L)$ compared to *AC-PG* $N^k(x)$ are lower. The size of the Aho–Corasick DFA depends on the underlying trie. While the maximum space complexity of the automaton in *AC-PG* $N^k(x|L)$ is $O(|N^k(x)| \cdot L)$, the automaton in *AC-PG* $N^k(x)$ has a maximum space bound of $O(|N^k(x)| \cdot (L + k))$. Additionally, we can say that $N^k(x|L) \subseteq N^k(x)$ since $N^k(x|L) = N^k(x) \cap \Sigma^L$ and an intersection is a subset of its parts. Thereby, Aho–Corasick DFAs for $N^k(x|L)$ are smaller than their counterparts for $N^k(x)$. Figure E.4 reflects this in the runtime ratios between the product graph using Ukkonens DFA and Aho–Corasick DFA, respectively. *Ukk-PG* $N^k(x|L)$ still performs better than *AC-PG* $N^k(x|L)$ due to smaller number of states and therefore smaller product graphs. The correctness evaluation for *Ukk-PG* $N^k(x|L)$ was evaluated on 114 904 strings (from 140 graphs) of which all matched the bottleneck costs computed with *Naive DP* $x|L$. Equivalently, all evaluated 68 465 strings for *AC-PG* $N^k(x|L)$ matched the bottleneck costs of *Naive DP* $x|L$. Figure E.1 and E.2 show the shares of strings covering graphs over the graph size $|E|$. Also here, the trends align with the results for Problem 2. Note, the difference ranges for x-axes as not all graphs were evaluated. The approximation results shown in Table E.1 indicate that while the errors are slightly larger compared to Table D.3, they remain fairly comparable and no under estimation of $B_x(t)$ occurs.

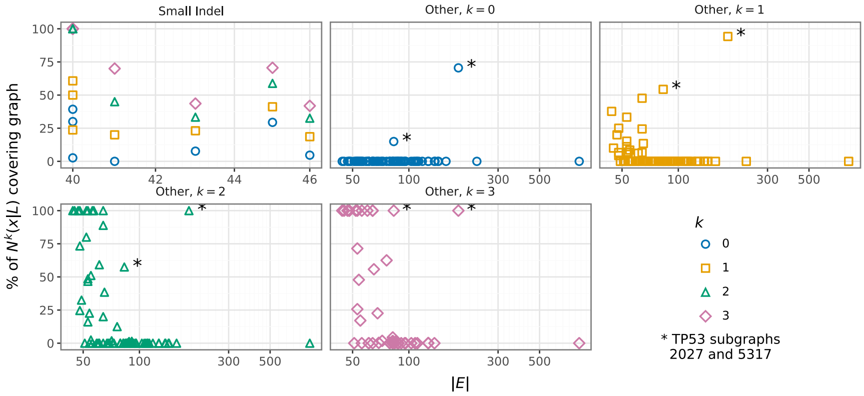


**Fig. E.1:** Results of *Ukk-PG* $N^k(x|L)$ for all three genes. With increasing graph size, fewer strings can cover the graphs. For many smaller graphs in *Other* there is a small percentage of candidate strings that can cover the graphs even with $k = 1$. Graphs marked with $*$ are larger near-linear graphs with 6 and 12 paths, respectively.

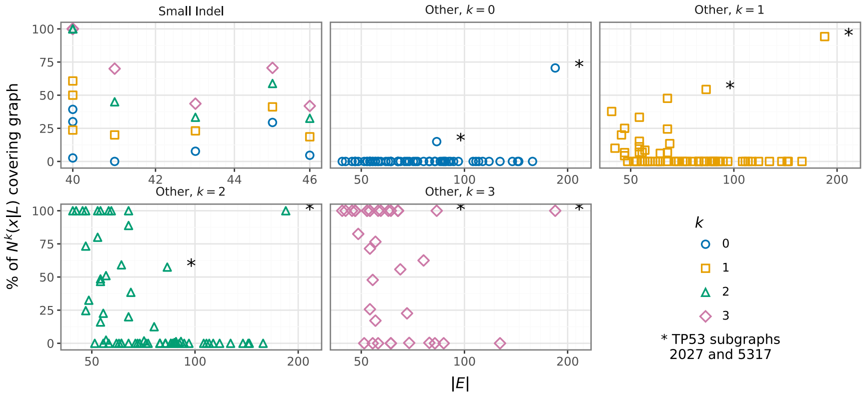


**Fig. E.2:** Results of *AC-PG* $N^k(x|L)$ for all three genes. With increasing graph size, fewer strings can cover the graphs. For many smaller graphs in *Other* there is a small percentage of candidate strings that can cover the graphs even with $k = 1$. Graphs marked with $*$ are larger near-linear graphs with 6 and 12 paths, respectively.

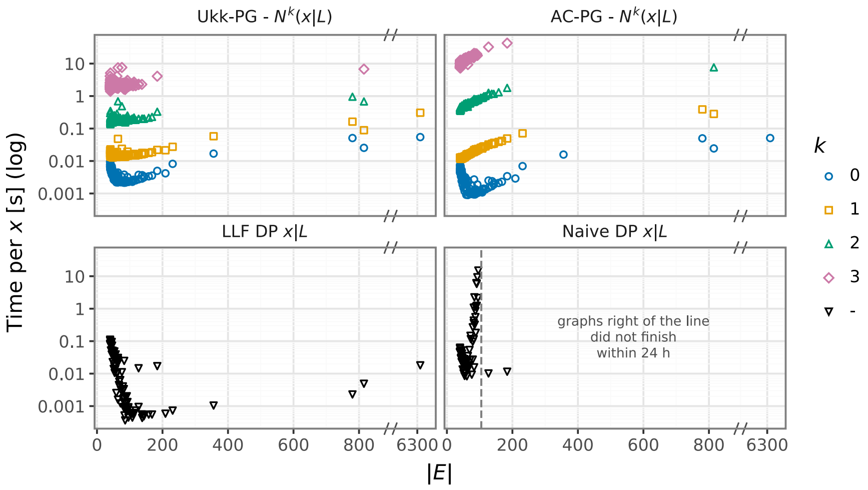


**Fig. E.3:** Comparison of runtime for *Naive DP* $x|L$, *AC-PG* $N^k(x|L)$ and *Ukk-PG* $N^k(x|L)$ over the number of nodes in the character-labeled graph representation for all subgraphs across three gene-specific pangenome graphs. This plot includes a reduced number of graphs as at the time of submissions nor all benchmarks could be completed.

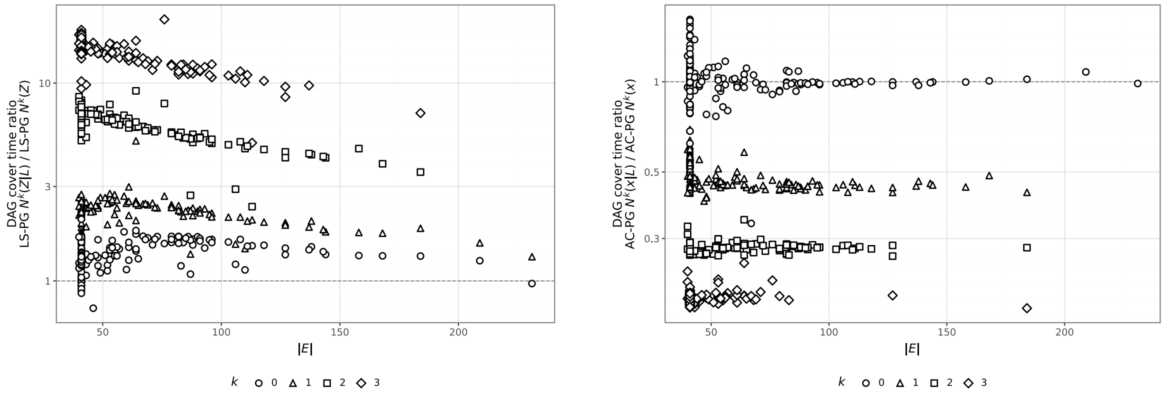


**Fig. E.4:** Ratio of runtime of *Ukk-PG* $N^k(x|L)$ over *Ukk-PG* $N^k(x)$ (left) and *AC-PG* $N^k(x|L)$ over *AC-PG* $N^k(x)$ (right), respectively.
